# Real-time targeted illumination in widefield microscopy achieves confocal quality neuronal images

**DOI:** 10.1101/2023.07.09.548132

**Authors:** Yao L. Wang, Jia Fan, Samuel H. Chung

**Affiliations:** Northeastern University, Department of Bioengineering, 360 Huntington Avenue, Boston, MA 02115

**Keywords:** neuronal imaging, widefield microscopy, target illumination, digital mirror device

## Abstract

Widefield fluorescence imaging has significant challenges in visualizing neuronal fibers near cell bodies. Specifically, out-of-focus and scattered light from the bright cellbody often obscures nearby dim fibers and degrades their signal-to-background ratio. Scanning techniques can solve this problem but are limited by reduced imaging speed and increased cost. We greatly reduce stray light by modulating the illumination intensity to different structures. We use a digital micromirror device in the illumination channel of a common widefield microscope and use real-time image processing to pattern the illumination. With the setup, we illuminate bright cell bodies with minimal light intensity, and illuminate in focus fiber-like structures with high light intensity to reveal weak signals. Thus, we minimize the background and enhance the visibility of fibers in the final image. This targeted illumination significantly improves fiber contrast while maintaining a fast-imaging speed and low cost. Using a targeted illumination setup in a widefield microscope, we demonstrate confocal quality imaging of complex neurons in live *C. elegans* and zebrafish larva, as well as in *in vitro* mice brain slice.

## Introduction

Widefield imaging is a conventional optical microscopy technique in which the entire field of view is illuminated and observed simultaneously [1-4]. While it offers the advantage of rapid image acquisition, it often suffers from lower contrast compared to scanning microscopy methods, such as confocal or multiphoton microscopy. This reduced contrast is due to limited optical sectioning and an inability to reject scattered light [5, 6]. In neuron imaging, this discrepancy is amplified by significant brightness differences between various structures under widefield illumination, which further undermines the contrast of finer structures. The neuron cellbody is generally larger in size and volume than axons and dendrites. For instance, the volume of a cellbody is around 113 μm^3^ with 6 μm in diameter, while the volume of an axon is 2.5 μm^3^ with 0.4 μm in diameter and 20 μm in length. Assuming identical fluorophore concentrations, the cellbody may emit roughly 45(113/2.5 = 45) times more photons than the axon due to the proportion of excitation photon counts to their volumes. Combined with the effect of the out-of-focus light, scattering light, and poor optical sectioning, the cellbody can often appear 20X – 30X brighter than axons and dendrites, thus obscuring nearby axons and dendrites (Fig. 1**Error! Reference source not found.**a). In contrast, scanning microscopy techniques, such as confocal or multiphoton microscopy, offer improved contrast by employing optical sectioning to eliminate out-of-focus light and reduce the impact of light scattering (Fig. 1**Error! Reference source not found.**b) [5, 7-9]. However, scanning microscopes are usually more complex and expensive, and often need to tradeoff between imaging speed and field of view (FOV), while widefield imaging preserves almost the same speed with a large FOV. Although scanning techniques provide more capability in imaging deeper along with higher signal-to-background ratio (SBR), widefield technique might be preferred when large FOV fast imaging is needed, or low-cost is important. Widefield imaging is even more preferred if their SBR can be improved.

**Figure 1.**
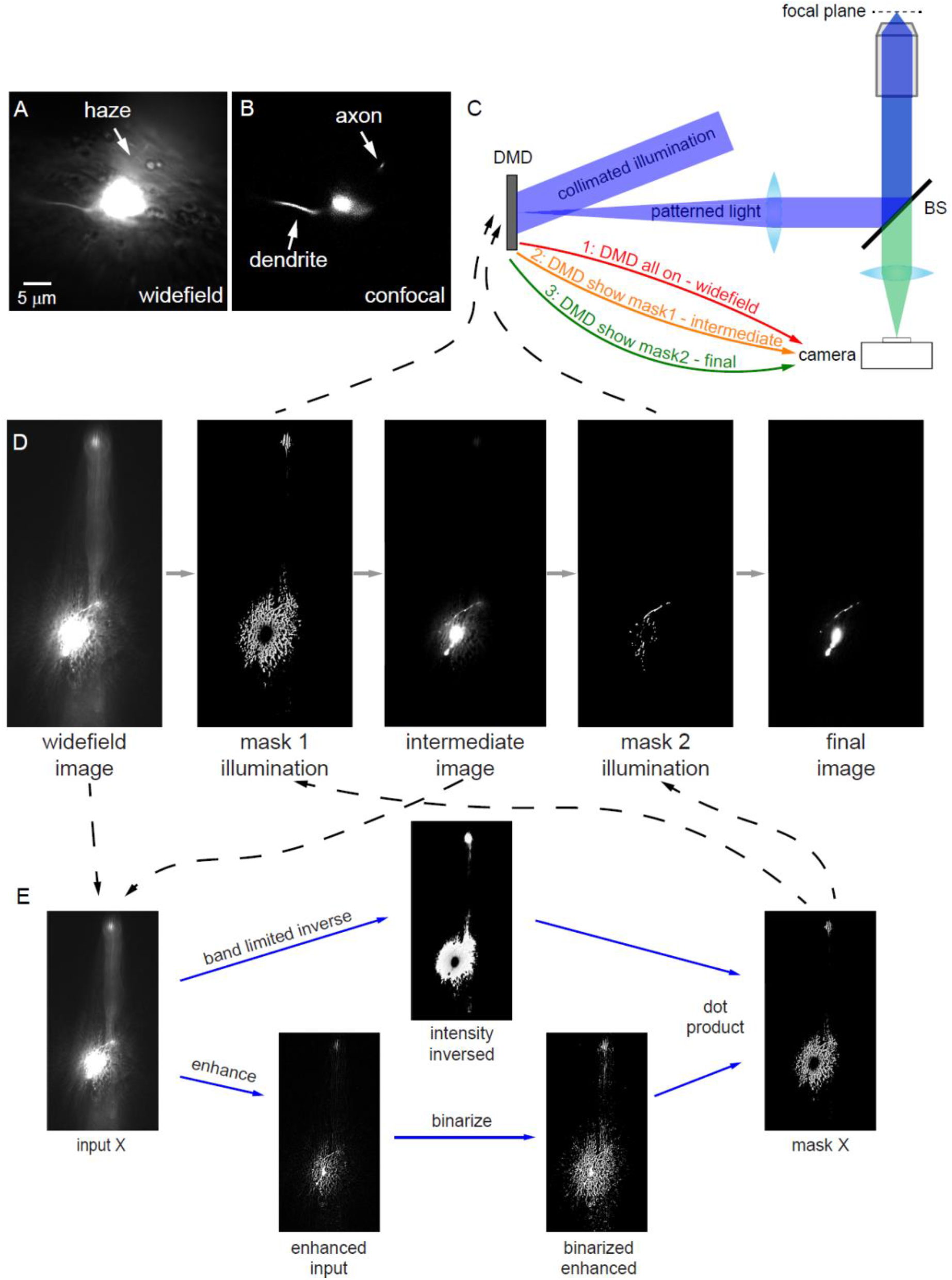
Principle of TIM. (**A**) The ASJ neuron of *C. elegans* imaged by widefield microscope. Dendrite and axon are relatively low contrast. (**B**) The ASJ neuron of *C. elegans* imaged by confocal microscope. Dendrite and axon are relatively high contrast. (**C**) Target Illumination Microscopy hardware setup. (**D**) Workflow of TIM with an example image of *C. elegans* ASJ neuron. (**E**) Flowchart of TIM mask generation. DMD: digital micromirror device. BS: beamsplitter.

In this study, we propose a novel approach termed Target Illumination Microscopy (TIM), which leverages targeted illumination to enhance the quality of neurite images in widefield microscopy [10, 11]. This technique illuminates only potential neuronal fibers using a simple Digital Mirror Device (DMD) on a widefield microscope platform (Fig. 1**Error! Reference source not found.**c), enhancing the signal contrast of these fiber structures and reducing the unwanted brightness of the cellbody and other background light, thereby improving image quality and contrast for the visualization of neurites and other intricate neuronal structures.

Successful TIM relies on generating accurate masks for neurite illumination. We hypothesize that no image processing method, including machine learning, can perfectly isolate only fiber-like structures without potential signal loss during mask generation. Therefore, we propose an iterative method that tolerates imperfect image processing (Fig. 1**Error! Reference source not found.**d). We first capture a widefield neuron image, then use image processing algorithms to generate an initial mask excluding most non-fiber-like structures. Using this initial mask, we pattern the illumination to obtain an improved image. From this improved image, we generate a second, more precise mask, resulting in a final image with minimal background light and significantly improved neurite contrast.

Using our TIM approach, we demonstrate improved neurite imaging in both *in vivo* and *in vitro* neuronal samples, including live *C. elegans*, zebrafish, and mouse brain slices. In all cases, the images acquired using TIM closely resemble those obtained using scanning techniques in terms of signal-to-background ratio (SBR). We believe that the development and application of TIM presents a significant step forward in the field of widefield imaging, offering a more accessible, cost-effective, and efficient method for high-quality neuronal imaging.

## Results

### Principle of TIM

During neuron imaging, selectively illuminating only in-focus fiber-like structures enhances the signal contrast of these fibers. This targeted illumination minimizes excitation photons on the cellbody and other background structures, thereby reducing their unwanted brightness. We propose a workflow illustrated in Figure 1d to achieve this targeted illumination imaging. We first capture a widefield neuron image and then employ image processing algorithms to generate a mask that ideally contains only in-focus fiber-like structures. However, due to the low-quality widefield image input to the image processing algorithms, it is often impossible to generate a perfect mask. With this imperfect mask, we pattern the illumination via a DMD [12-14] shown in Fig. 1**Error! Reference source not found.**c to obtain a new image. Because this mask eliminates light on most background structures, the resulting image presents much less background and cellbody haze compared to the original widefield image. We call this resulting image intermediate improved image. Then, we follow an iterative process to refine the image quality. We repeat the mask generation process using this intermediate improved image, allowing us to create a more precise mask. This new mask is then used to pattern illumination via the DMD, leading to a further enhanced image of neurites.

Fig. 1d illustrates the entire workflow of TIM, employing a straightforward *C. elegans* ASJ neuron image as an example. Initially, we capture a typical widefield image of the neuron. Based on this widefield image, we conduct image processing (discussed in later sections) to generate mask 1. With the aim of preserving all signals, we employ a lower threshold during image processing and segmentation to create mask 1, primarily to reduce the illumination of the background and cellbody. Patterned illumination through this mask results in an intermediate improved image. We then repeat the above procedures using this intermediate image but with a higher threshold in each step to create mask 2. Leveraging the superior quality of the intermediate image allows us to apply a higher threshold to generate mask 2 without risking the loss of neurite signals. This yields a much more precise mask 2. With this improved mask 2, the final neurite image is significantly enhanced.

Fig. 1e presents the general mask generation step, which encompasses the use of inverse widefield imaging to simulate Adaptive Illumination Microscopy [15-18], enhancing the widefield image to facilitate fiber-like structure segmentation, conducting fiber-like structure segmentation based on a specified threshold, and creating illumination mask through a combination of the inverse widefield and fiber-like structure segmentation. Each of these steps in TIM will be detailed in the subsequent sections.

### Inverse masks enhance dim signal contrast

Adaptive/Active Illumination Microscopy (AIM) is a technique utilized in scanning techniques. The AIM inverses the illumination laser power based on subject pixel brightness, and thus making excitation photons on bright structures much less and on dim structures much more [15-18]. This strategy greatly increases the imaging dynamic range and weak-signal sensitivity. By optimizing the illumination pattern, AIM also reduces photobleaching and phototoxicity, ensuring that live cells and delicate structures remain intact during imaging. We apply this AIM inverse illumination power idea in widefield imaging platforms as it has the potential to improve weak signals and suppress strong signals. The scanning AIM technique uses an electro-optic modulator together with a proportional-integral-derivative feedback circuit to pattern the light on each spot on the sample. In our widefield setup, we use the DMD to achieve similar functionality, which is to make illumination on each location different (Fig. 1**Error! Reference source not found.**c).

Leveraging the concept of inverse brightness, we first obtain a widefield image of the sample (Fig. 2a). Next, we invert its intensity within a specified range to create a mask (Fig. 2b). The upper limit of this range is typically set near the camera’s saturation value, given that the cellbody can exhibit brightness up to 20 times greater than the adjacent fibers. Conversely, the lower limit is generally set slightly above the dark count/noise floor of the CMOS (complementary metal–oxide– semiconductor) camera we employ [19]. Displaying this mask on the DMD results in patterned light illuminating the sample, thereby producing an enhanced neurite image (Fig. 2c). As fewer photons hit the cellbody, it appears dimmer than before, which subsequently reveals the proximate axon and enhances neurite contrast (Fig. 2cde).

**Figure 2.**
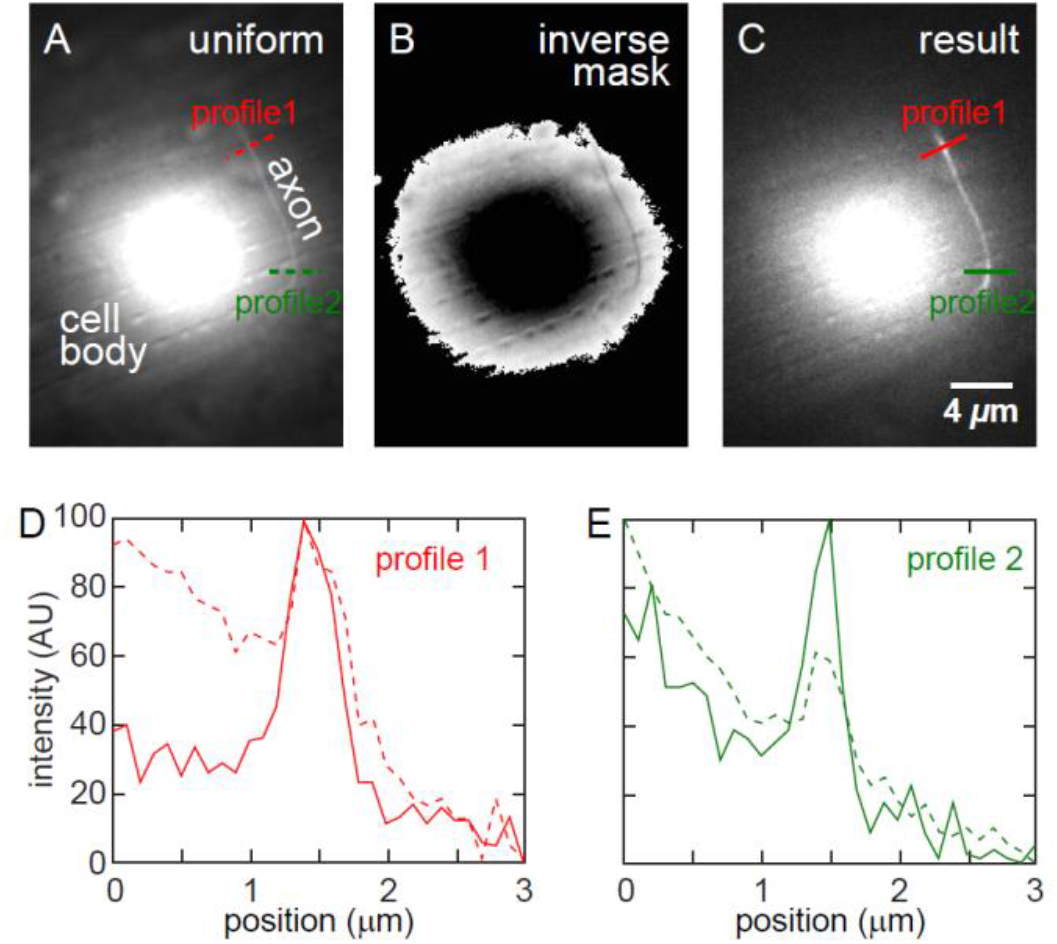
Improved fiber contrast with inversed illumination. (**A**) Widefield image of cellbody with uniform illumination. (**B**) Mask generated by inversing the original image in part A within an intensity range. This mask is displayed to the DMD, generating a patterned light on sample plane. (**C**) Image acquired by inverse mask illumination. Axon near to cellbody is relatively brighter as cellbody brightness is largely suppressed. (**D, E**) Normalized profiles of different locations of the axon. Red: location 1; green: location 2. Dash line: uniform illumination; Solid line: inversed illumination

In widefield imaging of neurons, the cellbody often appears much brighter than the surrounding neurites due to its larger volume and the light scattering properties of the tissue. In our widefield inverse illumination, despite attempts to minimize direct light on the cellbody within the in-focus plane, those out-of-focus planes and scattered light still contribute to the cellbody’s overall brightness. This level of brightness of the cellbody still preserves its haze effect, which overshadows the finer details of the nearby neurites, making them difficult to identify or visualize.

One reason we identified for the inability of inverse mask solely to improve widefield imaging is the presence of a substantial donut-shaped light surrounding the cellbody (Fig. 2b). This donut-shaped light interacts with the cellbody through out-of-focus illumination and scattering, leading to the emission of a large number of photons from the cellbody, which ultimately compromises the overall neurites image quality. Nevertheless, the inverse mask increases relative dimmer signal and suppresses relative brighter signal. Thus, we make the inverse mask one step in generating our TIM mask.

### Neuf significantly highlights fiber-like structures for high quality neurite segmentation

In image processing, a line detector is image filter employed to identify and extract linear features or structures within an image [20]. Fibers, to some extent, resemble lines. Thus, we use a variation of the line detector to highlight fiber-like structures in the original image. Since fibers generally have a relatively uniform diameter/thickness, we can first determine their thickness and then use this thickness to detect all line/fiber-like structures in an image. For example, most neurites under a combination of our microscope objective and camera possess a thickness of around 5 pixels, then we can design a line detector that specifically detects 5 pixels wide features in an image. For different objective and camera combinations, this number should be adjusted accordingly.

In the example shown in Fig. S1a, suppose there is a low neurite to be imaged. The line is a 5-pixel wide neurite with an intensity of 9, and the background should be uniformly 8. Due to an imperfect imaging system and inevitable noise, some parts of the neurite object appear as 8, and some background appears as 9 in the acquired image. One can imagine how challenging it is to precisely segment an exact 5-pixel wide line from this image using intensity-based thresholding. Since we know the neurite is approximately 5 pixels wide, we design a line detector spanning 5 pixels in width (Fig. S1a bottom left) and apply this detector/filter to the low contrast 5-pixel wide line image. This is a process of image filtering [21], similar to a bandpass filter in Fourier space, that passes some specific frequency signal while suppressing all other frequency signals. By convoluting the line detector with the original neurite image, we obtain a new image matrix (Fig. S1a right). Compared to the original image matrix on the top left, the resulting filtered image matrix shows a much higher contrast between the line signal and its background, and more importantly, the difficulty of segmenting this signal from the background is significantly reduced.

In real imaging, fiber-like structures mostly span through different directions than purely vertical or horizontal direction. To highlight them in all directions, we apply the line detector in both x and y directions and then combine them. The calculations we use are shown below. Note that, after one direction filtering, we set negative values to be zeros before inputting to the final Eq. 1. Negative values imply it is background with some brighter pixels nearby, and thus cannot be neurite signal pixel. If we keep this negative value to Eq .1, it will become a positive value because of the square.

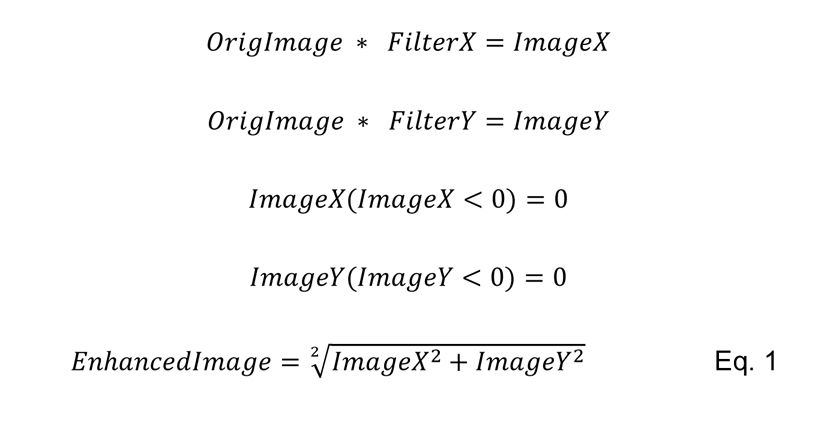

An example of using Eq. 1 to filter a widefield neuron image of *C. elegans* ASJ neuron is shown in Fig. S1b. All fiber-like structures are visibly enhanced, and all other signals/backgrounds are significantly suppressed.

The line detector/filter is designed specifically for neuron fibers and needs to be adjusted to accommodate different fiber thicknesses, whether it is due to differences in imaging set up, neuron differences within the same species, or neurons from different species. In order to differentiate this fiber filter from the typical line detector, which usually aims to detect single-pixel lines, we name our neuron fiber structure filter ‘Neuf’ (Neuron fiber filter). We use the name Neuf in the rest of this paper.

### Ratio image makes segmentation more reliable

Even with the Neuf enhancement, thresholding the fiber structure can still pose challenges due to the bright signals near the cell body maintaining very high values even after filtering. In Neuf filtering, the final filtered pixel values are “proportional” to the original values. As shown in Fig. 3a, in widefield imaging, due to scattered light from the very bright cellbody, fibers near the cellbody often appear brighter than fibers far away. Then, with the Neuf enhancement, enhanced fiber pixels near to cellbody still appear much brighter than distant fiber pixels (Fig. 3b and Fig. S2d), according to Eq. 1. This huge difference makes global thresholding based on this Neuf enhanced value risky. Moreover, different imaging conditions and samples may have very different overall brightness, necessitating adjusting the threshold for different images. Ideally, we want all desired signals to have similar values for easy segmentation.

**Figure 3.**
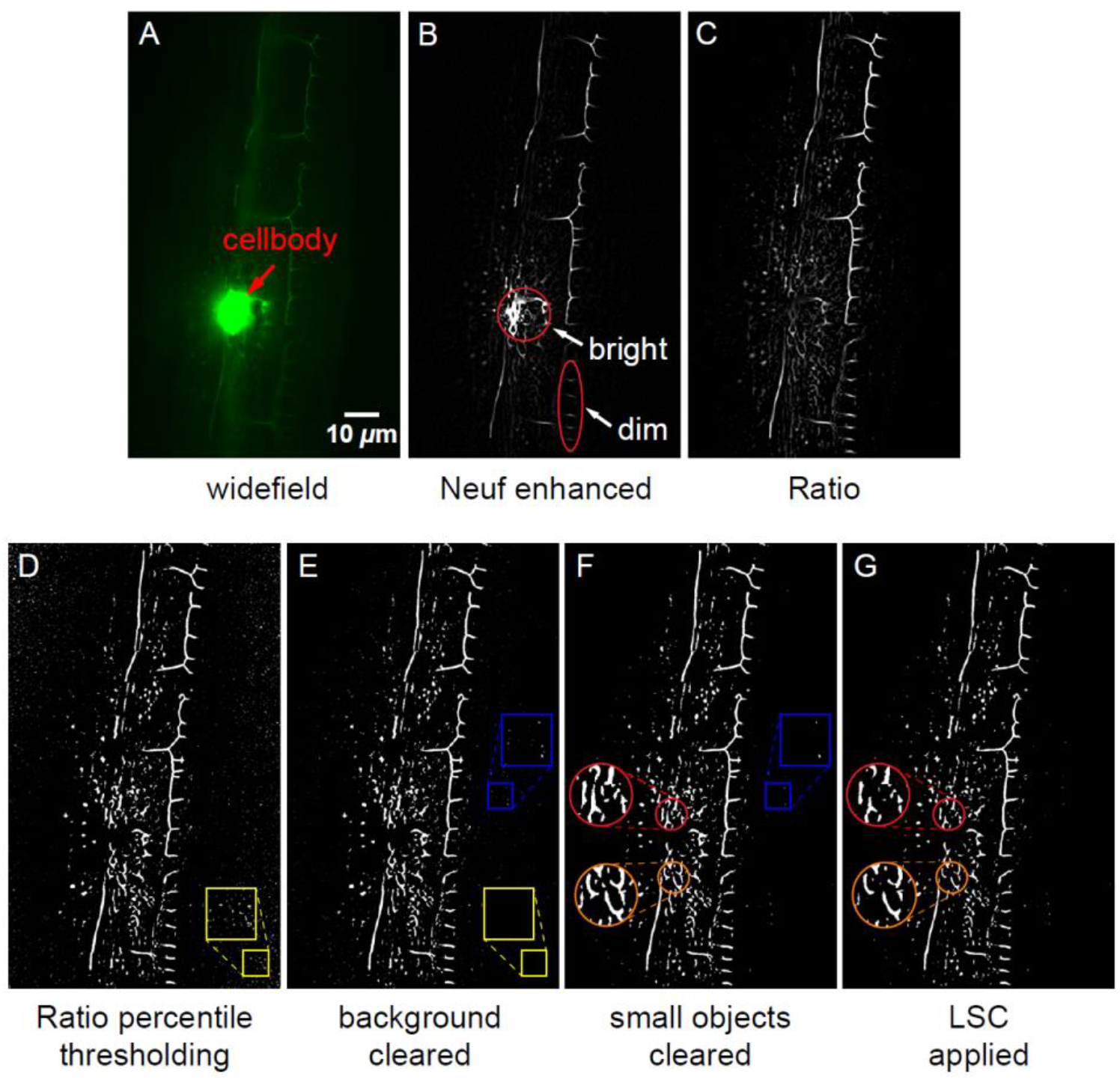
Fiber-like structure segmentation in TIM. (**A**) 2D plane of original widefield image of the PVD neuron in *C.* elegans. Fiber structures are much dimmer than the bright cellbody. (**B**) Neuf filtering original image in part A. Fiber structures are enhanced and cellbody is suppressed. (**C**) Ratio image, which is Neuf enhanced image in part B divided by the original image in part A. This kind of normalization further enhances contrast of all fiber like signal and suppress other background. (**D**) 96^th^ percentile thresholding of Ratio image. (**E**) Background cleared Ratio image. (**F**) Small objects cleared Ratio image. (**G**) Local sparsity constraint applied to moderate noise segmentation.

To resolve this problem, we use a Ratio operator to normalize fiber signal contrast. The Ratio is defined as ‘Neuf Enhanced value / Original value’. With this definition, both bright signal and dim signal are normalized to their original brightness, making this Ratio essentially indicates how similar the imaged structure is to fiber. In Ratio image, weak fiber-like signals stand out, and previously very bright signals become dimmer (Fig. 3c and Fig. S2e). The Ratio image shows how much the Neuf enhancement is of each pixel, while the Neuf simply enhances all fiber-like structures according to signals to surrounding contrast.

### Neuron sparsity enables using percentile for easy segmentation

Fiber signals in neuron images are typically sparse [22]. In *C. elegans* neuron imaging, if we divide a 3D z-stack image into submicron voxels, voxels that contain a real fiber occupy less than 1% of total voxels. In even denser samples like mice brain, fiber voxels occupy less than 2% of total voxels or still less than 1%. This is because all fibers are very thin, and neurons are usually selectively fluorescence labeled for imaging [23, 24]. This sparsity gives us a chance to use pixel value percentile to segment neurites, assuming all fiber-like pixels are in the high-end percentile.

Our Neuf-based Ratio image highlights all fiber pixels and suppresses all other pixels, which means that all fiber pixels values are at the high end of all pixels in that image. Suppose the fiber signal occupies 1% of the total pixels; then, of all pixels in that Ratio image, the top 1% or 2% of pixels should contain all fiber pixels. If we set the 96th percentile value as the threshold, the top 4% pixels will be designated as signal, and the rest 96% pixels as background. Fig.3d shows a 96th percentile segmentation based on the Ratio image. Although there are severe noises, all fiber structures are included in this segmentation. The sparsity of neurons defines this thresholding percentile, and thus we call this process sparsity-based thresholding.

An additional threshold must be applied to the Ratio image to obtain more precise segmentation, which is achieved through Neuf enhanced image thresholding and small connected object removal. Suppose a neuron image that contains only out-of-focus light and scattered light, and there is no fiber signal in this image, the percentile based thresholding will still produce a segmentation that contains 4% of all pixels, which are all false-positive signals. Looking back to the Neuf enhanced image, all out-of-focus and scattering light pixels have very low magnitudes, meaning a low threshold can easily remove them. For example, in a 16-bit (grayscale 0-65535) image, pixels with a gray value of 20 difference are usually within the same category (either signal or background), such as pixels with a value of 480 and pixels with a value of 500 are usually not two categories. If 20 is the difference, then the Neuf enhanced pixel value is 500 * 50 + 480*(-50) = 1000. This 1000 is the threshold that should be applied to the Neuf enhanced image to deal with a Z-stack imaging slice that contains no signal. In summary, we apply a relatively very low threshold to the Neuf enhanced image and combine this image (dot product matrix calculation) with the Ratio percentile segmented image to get the resulting image Fig. 3e.

After low magnitudes thresholding, there will be some very small objects with only 1-4 connected pixels on the segmentation, and those are so small and cannot originate from the signal. Thus, we also remove those very small objects from the segmentation in this step, as shown in Fig. 3f. Depending on different microscope objective and camera combinations, researchers should define the biggest object in their segmentation that should be removed.

We also compare the percentile based Ratio image thresholding to other potential fiber-like structure segmentation method and show the result in Fig. S3. In summary, for a widefield neuron image, the most widely used Otsu global thresholding [25, 26] seldom works (Fig. S3c); Sobel, Canny and LoG edge detection [26, 27] can extract many fiber-like features but fails to reject noise caused by tissue scattering, especially noise around cellbody (Fig. S3def). Local thresholding is a potential candidate to substitute the percentile based Ratio image thresholding, but it often fails to detect fiber features near the cellbody, as shown in Fig. S3ijk. Meanwhile, the sensitivity for local thresholding is hard to set automatically when the imaging subject changes.

### Local sparsity constraint confines highly scattering area segmentation

In certain instances, regions near the cellbody or other highly scattering tissues may produce exceedingly dense signals even after applying Ratio image percentile thresholding. To address this issue, we employ a local sparsity constraint (LSC) [28], which reduces the segmented signal density in these regions.

The LSC considers the unique characteristics of a surrounding area, names local, enabling the algorithm to selectively suppress dense signals that are likely to be artifacts or noise. This approach effectively maintains the relevant fiber signals’ integrity while minimizing the potential for false positives during the segmentation process.

To define ‘local’, we subdivide an image into numerous 50*50 pixel images, or 10x wide of a general fiber thickness. In every 50*50 region, we apply the LSC to moderate the noise segmentation. Considering that some regions intrinsically have denser signals than others, we set the LSC as 4 times the global sparsity. For example, if the global sparsity is 4%, names 96% percentile segmentation, then, in every local region, the local sparsity cannot surpass 16%, names 84% percentile segmentation based on the Ratio values. With this LSC, we make sure there is no segmented region that is too dense and looks bright, especially around the cellbody, which by itself fiber signal cannot surpass 16% sparsity, and this 16% sparsity is safe enough to include all possible fibers. Fig. 3fg shows the segmentation before LSC and after LSC. The local region signal/noise density is reduce by 1/3 and thus less noise will be produced in those regions during illumination.

### Directionality consideration combines low and high thresholding

While the Ratio image method substantially enhances all fiber-like structures in neuron images, there remains a small risk of losing valuable signal information during the percentile based segmentation process, particularly when employing an aggressively high sparsity threshold. Higher thresholds undoubtedly yield clearer segmentation results (Fig. S4a) but may inadvertently eliminate weaker, yet still relevant, signals. Conversely, lower thresholds tend to generate noisier segmentation outcomes but offer the advantage of maximizing the coverage and preservation of all signals, including those that are less prominent (Fig. S4b).

To address this tradeoff between clarity and signal preservation, our approach seeks to combine the benefits of both higher and lower sparsity thresholds to generate a clear yet robust segmentation (Fig. S4c). To link the high threshold segmentation and low threshold segmentation, we use double thresholding with directionality considerations [27]. The concept is borrowed from the Canny edge detection. First, when a pixel is higher than the high threshold, it is denoted as ‘1’. Second, if a pixel is higher than the low threshold and it is connected to a ‘1’ pixel with similar directionality, it will also be denoted as ‘1’. In this case, we maximize the possibility of covering all signals while eliminating most noisy background.

To determine whether two pixels have similar directionality, we first calculate their directionality with the range of -180° to 180° to produce a directionality map (Fig. S4d). Then, from their direction, we calculate their angle and check whether this angle is smaller than a defined ‘similar’ angle. For example, if we define angles less than 40° are similar angles, then, in Fig. S4e, the a° in the first quadrant is similar to (a + 40)° and (a - 40)°, and the b° in the second quadrant is similar to (-320 + b)° and (b-40)°. Note that this calculation and determination method differs from the most generally used direction angle determination method in the Canny edge detector [29]. In Canny, all directions are rounded to one of eight possible angles (namely 0, 45, 90, 135, 180, -45, - 90, or -135). In this case, a direction of 22° and a direction of 23° would be treated as having different directions, because the 22° will be rounded to 0 and the 23° rounded to 45. In our determination method, the angle between 22° and 23° is only 1°, which is within the 40° angle similarity and will be treated as having the same direction.

Based on our experience, considering directionality often proves to be redundant in most Ratio percentile-based threshold setups, as percentile thresholding is already robust against potential signal loss. We favor using a relatively lower sparsity threshold to prevent possible signal loss, as opposed to using this directionality consideration to recover potential signal loss during mask generation. Moreover, this directionality consideration is more computationally demanding compared to other processing steps. Therefore, we recommend users to employ this directionality consideration only when high fidelity of the image is crucial.

### Mask generation

By sequentially performing Neuf enhancement, Ratio image percentile thresholding, Local Sparsity Constraint (LSC), and directionality analysis (optional) (Fig. 4b), followed by a combination with the intensity inverse (Fig. 4c), we can generate a mask that encompasses all possible fiber-like structures and excludes very bright cellbody in neuron images (Fig. 4d). The rationale behind this specific sequence order is as follows:

**Figure 4.**
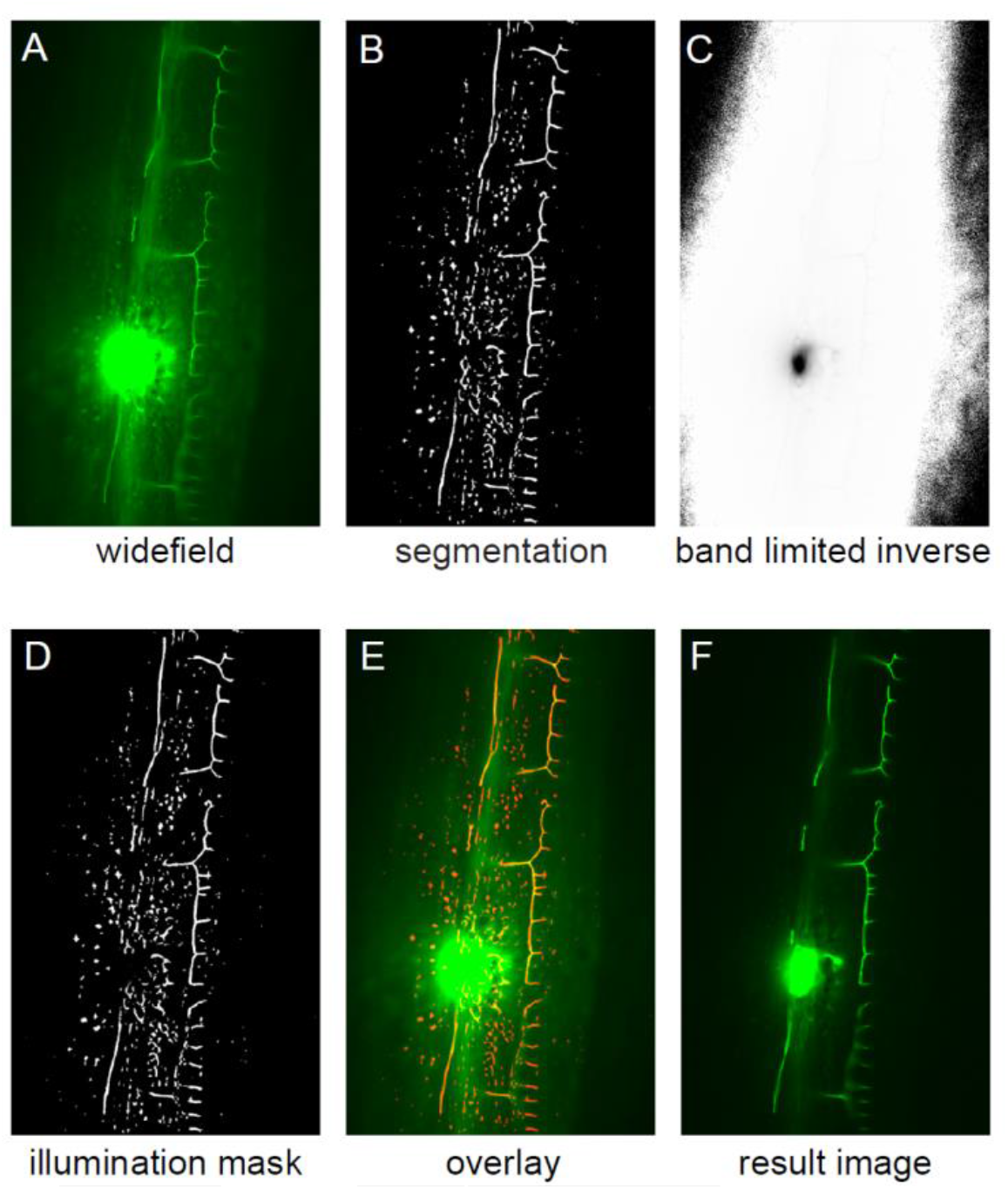
Mask generation. (**A**) 2D plane of original widefield image of the PVD neuron in *C.* elegans. (**B**) Segmentation of the widefield image based on Neuf, Ratio, LSC, and directionality in sequence. (**C**) Intensity inversed image of the original widefield image. (**D**) An illumination mask generated by the dot product of segmentation in part B and the inverse in part C. Note the cellbody area becomes black. (**E**) The overlay of generated mask on original widefield neuron image. The mask covers all possible fiber structures. (**F**) Image achieved with the generated illumination mask.

First, Neuf enhancement and Ratio image emphasizes fiber-like structures, providing a solid foundation for subsequent processing steps. Next, Ratio image percentile thresholding effectively separates the relevant fiber-like signals from the background noise depending on the fiber likelihood, further refining the image for segmentation. Then, the application of LSC aids in reducing noise density in regions near the cellbody or other highly scattering tissues to reduce potential artifacts. After that, directionality analysis is employed to identify the orientation of the fibers, providing valuable information about whether a pixel has more potential to be a signal pixel or a background pixel and reducing the risk of miss signals. Finally, combining the processed segmentation with the intensity inverse mask allows for highlighting weaker signals and suppressing strong signals during patterned illumination imaging. This sequence order has been carefully designed to maximize the clarity and reliability of the mask generated based on widefield images. A generated mask is overlayed with the original widefield neuron image in Fig. 4e to show the reliability of this mask generation method. With the mask being displayed in DMD and mask patterned light exciting the sample, an improved image is achieved (Fig. 4f).

### TIM full workflow

With the generated mask, we can get an improved but still imperfect image, as there is still a large amount of haze around the cellbody and out-of-focus light near many fibers. Simply increasing the threshold applied to each step in the mask generation process can produce a clearer image but suffers from the risk of losing signals. We believe it is impossible to generate a perfect mask based on a bad input image (widefield image in our case), even with more advanced image processing or computer vision algorithms.

As shown in Fig. 1d, we iterate the mask generation based on the intermediate improved image acquired with the first generated mask and then follow the previous Neuf, Ratio, LSA, and directionality to generate a second mask. Because the second mask is generated based on the improved image, the mask can be more precise and robust to signal loss with a higher threshold during segmentation (like change the Ratio percentile threshold from 96% to 98%).

Note that the intensity inverse is not strictly executed in the second mask generation. We perform every other segmentation procedure (Neuf, Ratio, LSC, and directionality) based on the intermediate improved image, but we reuse the intensity inverse image based on the original widefield image. Also, this inverse image is binarized instead of keeping its original bit-depth so that the final mask is a binary mask instead of a grayscale mask.

In total, we generate two masks to obtain the final TIM image. The iterative process can be repeated multiple times to refine the segmentation results further. However, practical considerations limit the number of iterations, such as imaging speed and the potential for photobleaching due to prolonged exposure to light. Balancing these factors, we have chosen to stop at the second mask, which provides an optimal compromise between image quality, imaging speed, and photobleaching/phototoxicity.

Moreover, the TIM method’s iterative nature can be adapted to various imaging modalities and applications, offering a versatile solution for situations where image quality and segmentation accuracy are paramount. This flexibility makes TIM a valuable tool for researchers across various scientific disciplines.

### TIM improves widefield neuron image step by step

We next perform a real experiment to show how the TIM improves neurites contrast in widefield imaging step by step.

For reference, we take a grid scanned image of the PVD neuron in *C.elegans* (Fig. 5a), as used in previous sections. Grid scanning is an established technique that is akin to spinning confocal but slightly inferior to point scan confocal [4, 30, 31].

**Figure 5.**
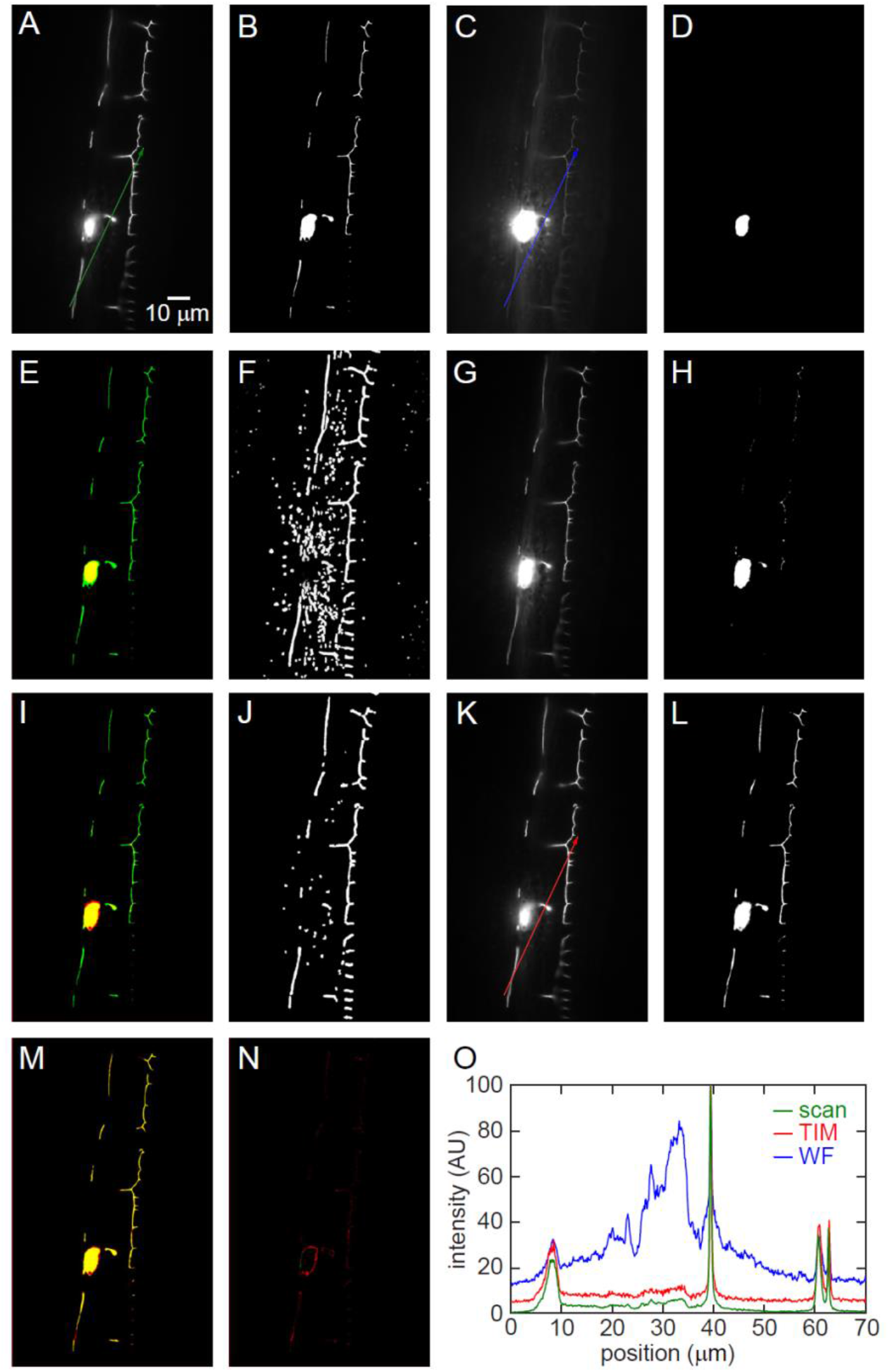
TIM improves imaging step by step. (**A**) Grid scan image of the PVD neuron in *C. elegans*. (**B**) Segmentation of the grid scan image using Otsu method. We use this segmentation as reference. (**C**) Original widefield image of the neuron. (**D**) Segmentation of the widefield image using Otsu method. (**E**) Overlay of the widefield segmentation (red) and grid scan segmentation (green). Note that the superposition of red and green is yellow. (**F**) Mask 1 generated based on the image in part C. (**G**) Mask 1 illumination acquired intermediate improved neuron image. (**H**) Segmentation of the intermediate image using Otsu method. (**I**) Overlay of the intermediate segmentation (red) and grid scan segmentation (green). (**J**) Mask 2 generated based on the image in part G. (**K**) Mask 2 illumination acquired further improved neuron image. This image serves as the TIM final image. (**L**) Segmentation of the TIM final image using Otsu method. (**M**) Overlay of the TIM image segmentation (red) and Grid scan segmentation (green). Almost all foregrounds are yellow. (**N**) The difference between TIM image segmentation and grid scan segmentation, colored with red. (**O**) Profile analysis based on lines in parts A, C, K.

To compare whether one neuron image is better than another, several methods are available, including profile analysis at specific locations, visual inspection, and segmentation results comparison. The profile analysis can only show the comparison in specific locations but not the whole image. Visual inspection can often be biased, and it is challenging to compare subtle contents. Here, we use the segmentation results comparison method to compare neuron images, and the segmentation results serve as an indicator of how good the neuron image SBR is. We use the mostly accepted k-means/Otsu algorithm among many segmentation techniques for this task. This algorithm separates all pixels in an image into two classes, foreground, and background. The threshold is determined by minimizing intra-class intensity variance, or equivalently, by maximizing inter-class variance [25, 26]. If all signals and background pixels are distinct from each other in an image, then the k-means/Otsu method can perfectly segment the image into foreground signal and background.

Because the grid scanning produces a reliable image, the Otsu segmentation result is reasonably decent and serves as a segmentation reference (Fig. 5b). As shown in Fig. 5cd, Otsu segmentation based on the original widefield image treats only the cellbody as foreground signals. The overlay of grid scan segmentation (green color in Fig. 5e) and widefield segmentation (red color in Fig. 5e) shows huge differences.

With the original widefield image, we were able to generate mask 1, which still contains severe noisy background (Fig. 5f). Nevertheless, with mask 1, we achieved an intermediate improved image (Fig. 5g). The improved image demonstrates a dimmer cellbody compared to the original widefield image, and the brightness of the relative dim fiber structures are significantly improved with more uniform grayscales. Based on this image, the same Otsu algorithm produces a better segmentation, shown in Fig. 5h, but still not comparable to grid scan image segmentation (Fig. 5i). A mask 2 generated based on this intermediate improved image (Fig. 5j) achieves a further improved image (Fig. 5k). The Otsu algorithm produces a new segmentation, shown in Fig. 5l, almost identical to the grid scan image segmentation (Fig. 5m). The segmentation results difference between this TIM final image and grid scan image is minor (Fig. 5n). Profile analysis at specific locations confirms that TIM provides better SBR than widefield and is similar to Grid scan (Fig. 5o).

### TIM *in vitro* imaging results

We first performed *in vitro* imaging on mice brain slices to confirm the TIM imaging capacity on neurons. The slices contained multiple neurons cortex region, labeled with green fluorescent antibodies. The brain slices were prepared using the method described in Ref. [32], and the thickness is around 40 μm.

In our TIM workflow, we need to set a few parameters for Z-stack imaging, including illumination intensity, exposure time, Z range, and the sparsity of the sample. The only TIM-specific parameter is the sparsity, and we set it to 4% for mice brain slice imaging, as they appear dense.

Brain slices are known to be highly scattering and tend to contain a substantial amount of out-of-focus light [33], primarily due to the density of neurons in all three dimensions (XYZ). Consequently, the original widefield images of brain slices often exhibit a very bright background and low contrast neurites, making it challenging to distinguish individual fibers and their intricate structures (Fig. 6a). This issue arises from the lack of optical sectioning capacity and the inability to reject scattered light in widefield microscopy, which results in some neurites becoming indistinguishable from their background. In contrast, our TIM imaging method significantly improves image quality by providing optical sectioning capabilities, as its mask illuminates only in-focus structures, and light to out-of-focus structure is quickly diverged, as discussed in our previous research [4]. This approach helps to mitigate the effects of out-of-focus light and scattering, enabling the visualization of much clearer and higher contrast neurite images (Fig. 6b).

**Figure 6.**
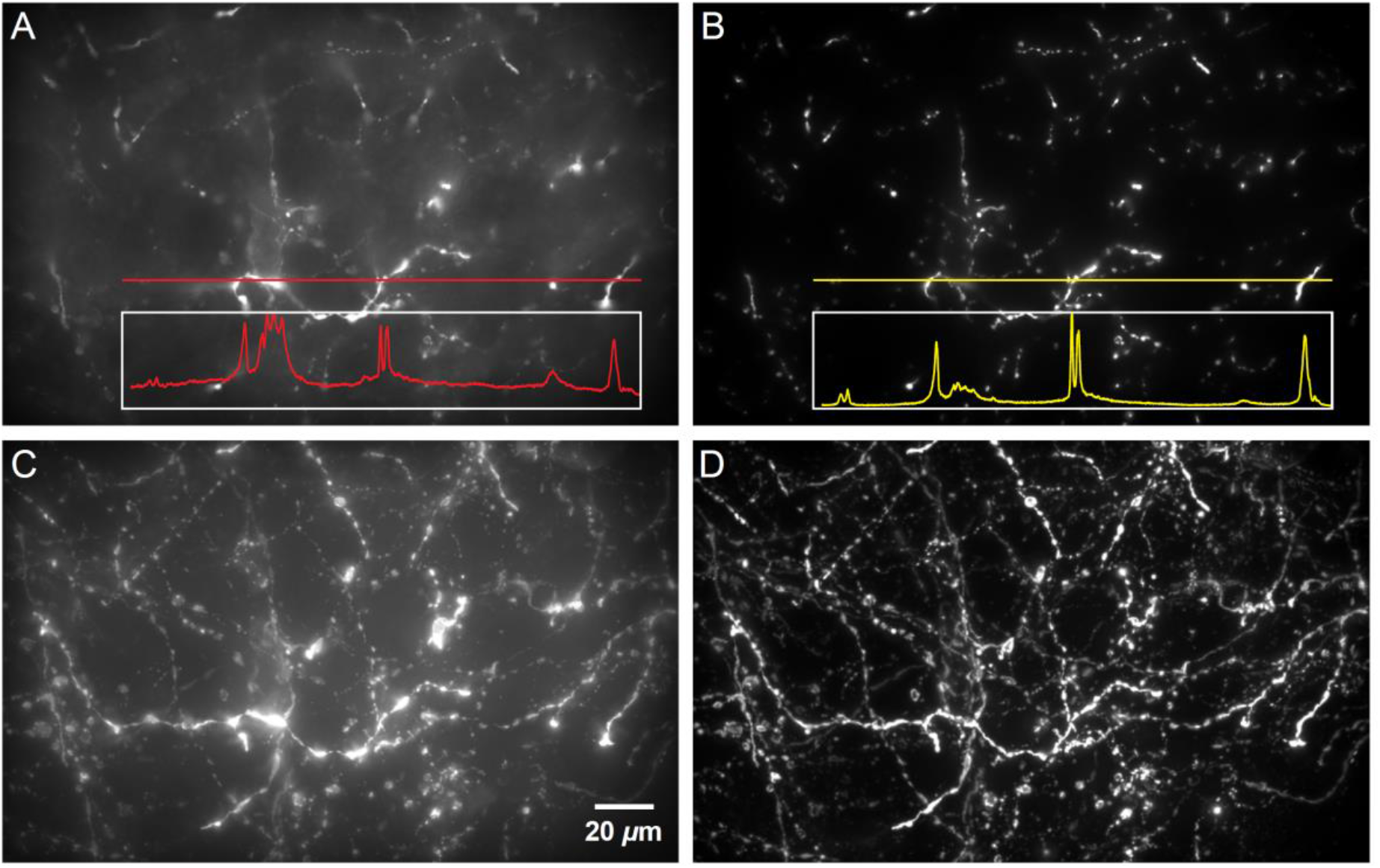
TIM imaging of mice brain slice. (**A**) Original widefield imaging of a 2D plane of a green fluorescent antibody labeled mice brain cortex slice. Red profile (normalized to 0-1) shows low neurites contrast. (**B**) TIM imaging of the same cortex slice. Yellow profile (normalized to 0-1) shows high neurites contrast and reduced out-of-focus light. (**C**) Max projection of the 40mm thick brain slice widefield Z-stack image. (**D**) Max projection of the 40 mm thick brain slice TIM Z-stack image.

We executed widefield and TIM imaging throughout the entire 40 μm brain slice, utilizing 0.5 µm z-steps to achieve a Z-stack image of the sample. The maximum intensity projection of the widefield image is shown in Fig. 6c. Due to the inherent limitations of widefield imaging, such as poor optical sectioning and high background noise, many neurites appear to be buried in the background, making it difficult to distinguish them from the surrounding structures. This lack of clarity hampers the accurate representation and analysis of the neural networks within the sample. In contrast, the TIM Z-stack maximum intensity projection, shown in Fig. 6d, reveals a significantly greater number of neurite structures with a substantially lower background. The enhanced clarity and contrast achieved through the TIM method allow for more accurate visualization of the intricate neuronal networks and structures, facilitating a better understanding of the underlying biological processes.

### TIM *in vivo* imaging results

Zebrafish is a complex model organism widely used in various fields of biological research [34-37]. Its transparent embryo, rapid development, and genetic similarities to humans have made zebrafish a popular model for studying vertebrate development, organ function, disease modeling, and drug discovery (Fig. S5a). Since the zebrafish has increased size, less transparent tissue, deeper depth of neuron, and more movement during immobilization, the imaging experiment on zebrafish is more challenging, and most researchers perform high resolution zebrafish neuron imaging only with advanced microscope platform like spinning confocal and two photon microscopy [38-41]. Widefield imaging often fails to resolve the fine structure of zebrafish neurites.

To confirm the TIM imaging capacity on *in vivo* neurons, we further performed TIM on a *l2b:Gal4 UAS:Dendra* line zebrafish. In Fig. 7, we showcase a comparison between widefield imaging and TIM imaging in the hindbrain region of the fish, focusing on the trigeminal neuron (Fig. S5b).

**Figure 7.**
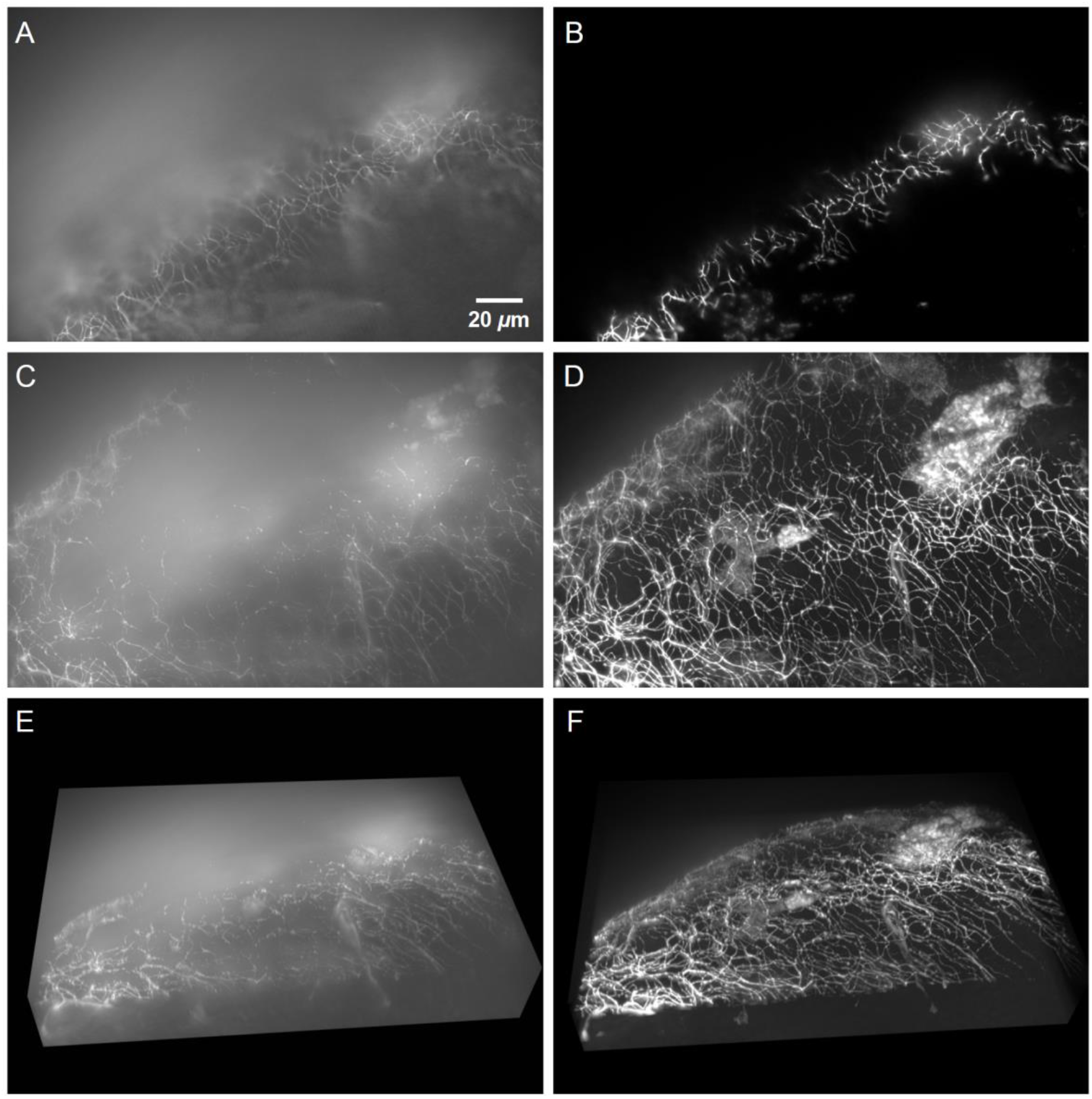
TIM imaging of live zebrafish. (**A**) 2D plane of GFP labeled trigeminal neuron, image taken with 60X widefield epifluorescence imaging modality. Massive background light are around both neurites and other areas. (**B**) 2D plane of GFP labeled trigeminal neuron, image taken with 60x TIM imaging modality. (**C**) Max intensity projection of 45 μm Z-stack of GFP labeled trigeminal neuron under widefield imaging. (**D**) Max intensity projection of 45 μm Z-stack of GFP labeled trigeminal neuron under TIM imaging. (**E**) 3D volumetric rendering of widefield neuron Z-stack image. (**F**) 3D volumetric rendering of TIM neuron Z-stack image.

Lack of optical sectioning capability, the widefield image contains massive and bright background (Fig. 7a). In contrast, the TIM method significantly reduces background noise near neurites (Fig. 7b). The enhanced clarity offered by the TIM method is further emphasized in regions away from neurites. Due to the minimal illumination after mask patterning, these areas appear purely dark, serving as an ideal background that highlights the neural structures. The max projection comparison in Fig. 7cd and the 3D volumetric rendering of the neuron images in Fig. 7ef further demonstrate our TIM technique’s enhanced neurite imaging capabilities.

## Discussion

To conclude, our TIM presents an economical and pioneering solution for neurite imaging that addresses some key drawbacks of traditional widefield microscopy. TIM’s optical sectioning capacity minimizes out-of-focus light, enhancing image clarity.

There are four important strengths of TIM. First, its affordability. The required DMD or similar Spatial Light Modulator (SLM) attachment to a standard widefield microscope is a relatively economical addition compared to more sophisticated platforms such as spinning confocal and two-photon microscopy. This cost-effectiveness broadens the accessibility of the TIM technique to a wider range of researchers and institutions. Secondly, it is easy to achieve fast imaging speed. For a 2D slice imaging, only three camera-acquired images are required, whereas scanning techniques necessitate millions of voxels to obtain a comparable image. The TIM relaxes the need to scarify its FOV for imaging speed or vice versa (to some degree), while scanning techniques always need to balance its imaging speed and FOV, as they usually persist the same imaging throughput [10, 42]. Third, TIM reduces photobleaching. The initial exposure uses low-intensity light, and subsequent exposures selectively target only neurite structures, thereby reducing scattered photons to neurites. In contrast, scanning techniques indiscriminately illuminate all regions [43], increasing photobleaching and potentially damaging the sample. Fourth, TIM can be used for longevity fast neurite activity tracking. Once a TIM mask is generated, similar to previously demonstrated high speed neuron activity tracking[11, 44], we can use the same mask for the remainder of the imaging, with speed limitations dictated only by the camera and refocusing unit speed. This strategy allows faster imaging than scanning techniques, although the mask may need small adjustment to account for animal motion effect. This TIM based neurite activity tracking also potentially reduces the photobleaching and thermal effect, compared to scanning techniques which illuminate all regions regardless of target structure presence.

The idea of iteration behind TIM can also serve as a verification method for machine learning enhanced images or segmentation. In machine learning for neuron image enhancement, they generate new or enhanced neuron slices based on fewer Z-slices [45]. However, human inspection is the only way to verify whether their generated new slices or enhanced image is trustworthy. With the TIM procedure, they can generate a mask based on their slice and use it to project patterned illumination to the sample. If the sample returns with signal on all locations, that means their generated or enhanced image is trustworthy, or vice versa.

The TIM technique offers several distinct advantages over traditional widefield imaging and scanning techniques. By providing higher contrast imaging than widefield and faster image acquisition than scanning, TIM can lead to more efficient data collection and preservation of delicate samples.

In total, our TIM requires three camera acquisitions time, three DMD response times, and two mask generations time. The combined duration of these processes often exceeds one second, which may be considered a limitation of our current implementation. However, several potential ways exist to improve the efficiency and speed of the TIM technique. For instance, upgrading our current slow projector-extracted DMD to a scientific DMD with a faster response time can significantly reduce the time required for pattern display. Furthermore, optimizing the mask generation algorithm and implementing GPU computing using CUDA and OpenCV can accelerate mask generation, streamlining the entire process [46, 47]. With these improvements, the TIM method could become an even more powerful tool for neuronal imaging, maintaining its advantages in terms of image quality, optical sectioning, and reduced photobleaching, while significantly increasing imaging speed.

## Materials and Methods

### Microscopy setup

All optics components are attached to a commercial Nikon Ti2-E inverted microscope for imaging convenience. All fluorescence images were acquired by a 1.4 NA, 60x oil immersion objective and a Hamamatsu ORCA-Fusion BT sCMOS camera.

### DMD setup

Similar to a prior studies [4, 48], we removed a DMD from a digital projector (Vivitek DW814). The DMD consists of 1280×800 micro mirror elements/pixels with 0.65-inch diagonal size. The DMD is placed in a conjugate plane to the imaging plane, which is formed by placing a tube lens (Thorlabs TTL200) on the back port of an inverted microscope (Nikon Ti2). The DMD and tube lens are aligned by eye. We use a 5-axis stage (Thorlabs PY005) and a Kinematic 1” Optic Mount (Thorlabs KS1) to hold and align the DMD. We use Matlab 2022a (MathWorks) to display mask/illumination pattern on the DMD. The code is available at Github (https://github.com/wormneurolab/DMD-TIMconfocal).

### DMD alignment

Utilizing the center row and column elements of the DMD, we project an image of a cross onto a thin fluorescent film. We adjust the DMD position and orientation using the 5-axis stage so that the cross is in focus and centered on the camera array. The thin fluorescent film is made by drawing a 3-mm diameter spot on a microscope glass slide. We cover the spot with a coverslip and applied pressure, generating a thin film between the coverslip and slide. We tape two edges of the thin film slide and allowed it to dry at room temperature for 4 hours.

### Illumination and imaging

We use an LED light source (Lumencor SOLA light engine) through a liquid guide as fluorescence illumination. After collimation (Thorlabs SM1U25-A), the illumination is shined to the DMD. We use MATLAB to treat the DMD as the second monitor of our computer and project the generated mask to the DMD to pattern the illumination. The exposure time depends on intensity of the light source, transmission of the DMD, strength of fluorescent labelling, and sensitivity of the camera. We use 0.05-s exposure time for widefield and DMD-enabled imaging. Utilizing our setup and samples, we acquired all the raw data for a 2D DMD image in ∼ 1 s. This extended time was primarily due to the slow responding time of the projector extracted DMD and our Matlab image processing algorithm, especially the ‘directionality consideration’ step.

### Animal preparation

The *C. elegans* are cultivated using established protocols on Bacto agar plates [49] at 20 °C. When imaging, worms are in their young adult stage.

The mice brain slice is prepared using the method described in Ref. [32], and the thickness is around 40 mm.

The zebrafish larvae used in this research is a *l2b:Gal4 UAS:Dendra* line zebrafish. This line uses GFP to label the retinal ganglion cells and the trigeminal nerves [50, 51]. The zebrafish and zebrafish larvae are cultured under established protocols [52]. When imaging, the zebrafish larvae are around one week old, after hatching.

### Imaging of Animals

The *C. elegans* is immobilized by sodium azide in 2% agarose pad on microscope glass slide and covered with coverslip. The sodium azide paralyzes the worm so it has zero movement during imaging.

The Zebrafish larvae were mounted on a microscope slide 2% low-melting-point agarose (Sigma) and covered with coverslip. The agarose was sufficient to restrain movements of the larvae and we didn’t apply anesthesia [53]. Larvae can be maintained in this slide for imaging sessions lasting more than 4 hour, and they can develop normally after the imaging and take off from the slide.

## Supplemental Data

**Figure S1. Neuf for highlight fiber-like structures.** (**A**) A line image with low contrast is convoluted to a filter that is designed specifically with the line’s width. After convolution, the result line image demonstrates a much higher contrast against its background. (**B**) Conventional widefield image of ASJ neuron is convoluted and processed by the Neuf filter in both XY directions. The resulting image highlights all in-focus fiber-like structures while suppressing all other signals.

**Figure S2. Different percentile segmentation on Neuf image and Ratio image.** (**A**) Widefield image of a PVD neuron in *C. elegans.* (**B**) Widefield image of a PVD neuron in *C. elegans.* with bright light. (**C**) Widefield image of a PVD neuron in *C. elegans* with dim light. (**D**) Neuf enhanced images with different percentages. (**E**) Ratio images with different percentages.

**Figure S3. Different image processing methods for fiber detection.** (**A**) Widefield image of a PVD neuron in *C. elegans.* (**B**) Segmentation of original image using 98^th^ percentile (top 2%) thresholding on Ratio image. (**C**) Segmentation using Otsu global thresholding of original image. (**D**) Segmentation using Sobel edge detection on original image. (**E**) Segmentation using Canny edge detection on original. (**F**) Segmentation using LoG edge detection on original image. (**G**-**K**) Segmentation using local thresholding on original image, with different sensitivity.

**Figure S4. Directionality map connecting with relatively weaker signal.** (**A**) Apply a relatively high threshold on the Ratio image to segment. (**B**) Apply a relatively low threshold on the Ratio image to segment. It shows more noise and potential fiber structures. (**C**) A segmentation based on considering both high threshold, low threshold, and the directionality similarity. Note the left red circle shows that this combination connects fiber-like structures in high thresholding by directionality similarity connection, and the right circles show that this combination doesn’t significantly expose more noise like in low thresholding. (**D**) The directionality map is calculated based on Neuf magnitudes and ‘atan2’ function in Matlab. Connected fibers appear to have similar directions. (**E**) The directionality angle judgment in our algorithm. Directions with a small angle difference are treated as having similar directionality, instead of dividing them into different subareas in Canny edge detection.

**Figure S5. Zebrafish imaging area.** (**A**) Live zebrafish fixed to agarose pad to mechanically constrain its movement. Image taken with 2X brightfield imaging modality. (**B**) GFP imaging region in red rectangle. Image taken with 20X brightfield imaging modality.

## Notes

### Competing Interest Statement

The authors have declared no competing interest.

