## Supplemental figures for "Real-time targeted illumination in widefield microscopy achieves confocal quality neuronal images"

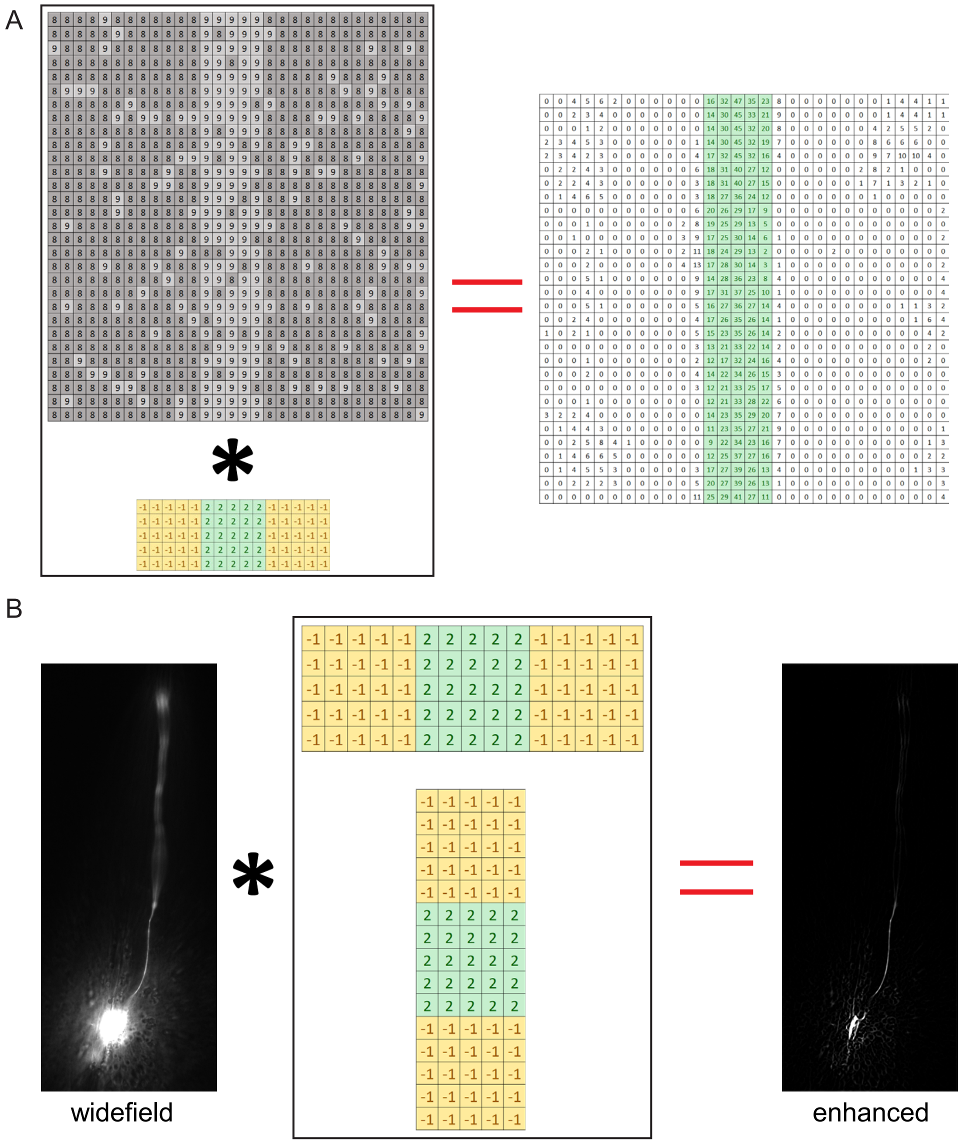


**Fig. S1. Neuf for highlight fiber-like structures**

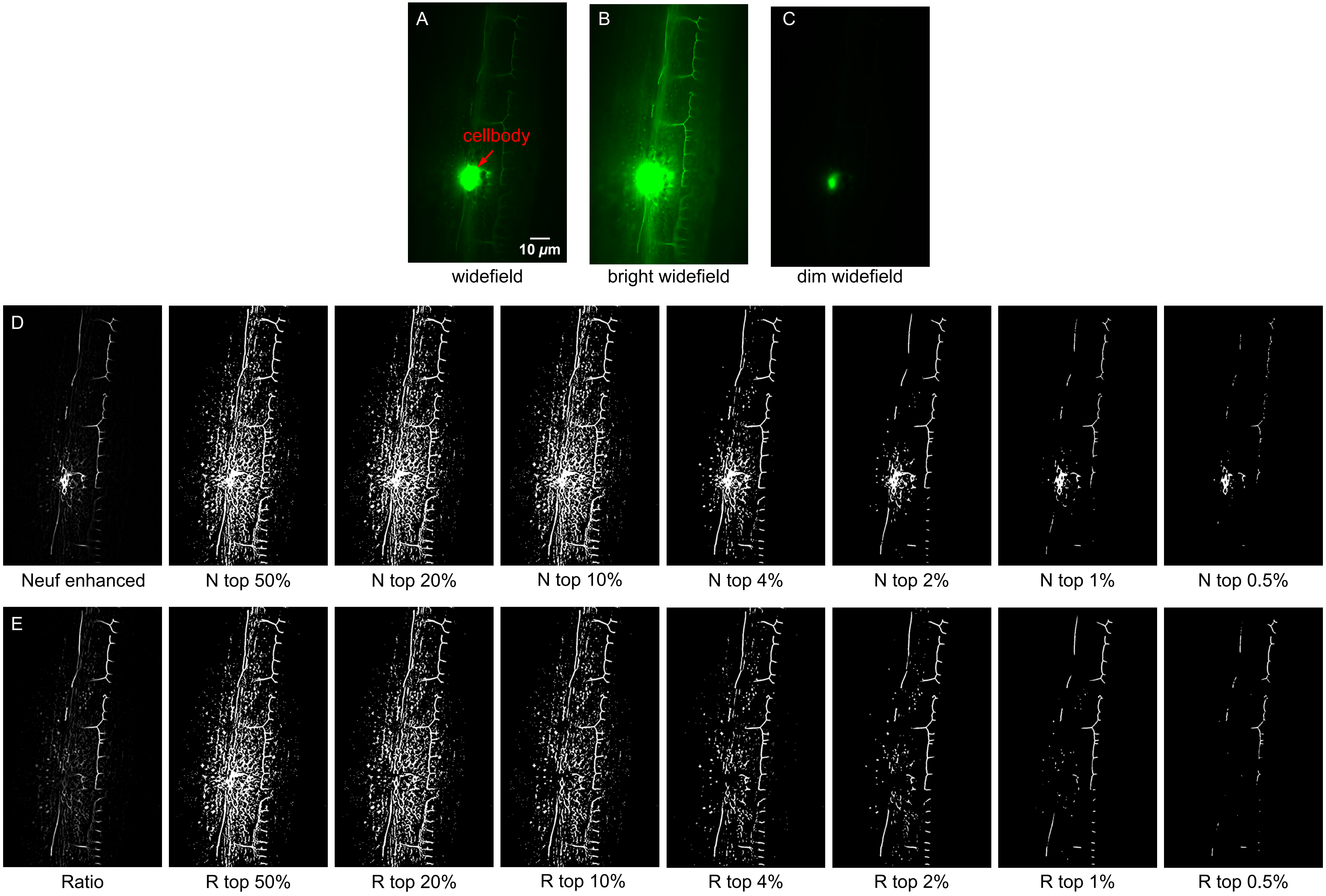


**Fig. S2. Different percentile segmentation on Neuf image and Ratio image**

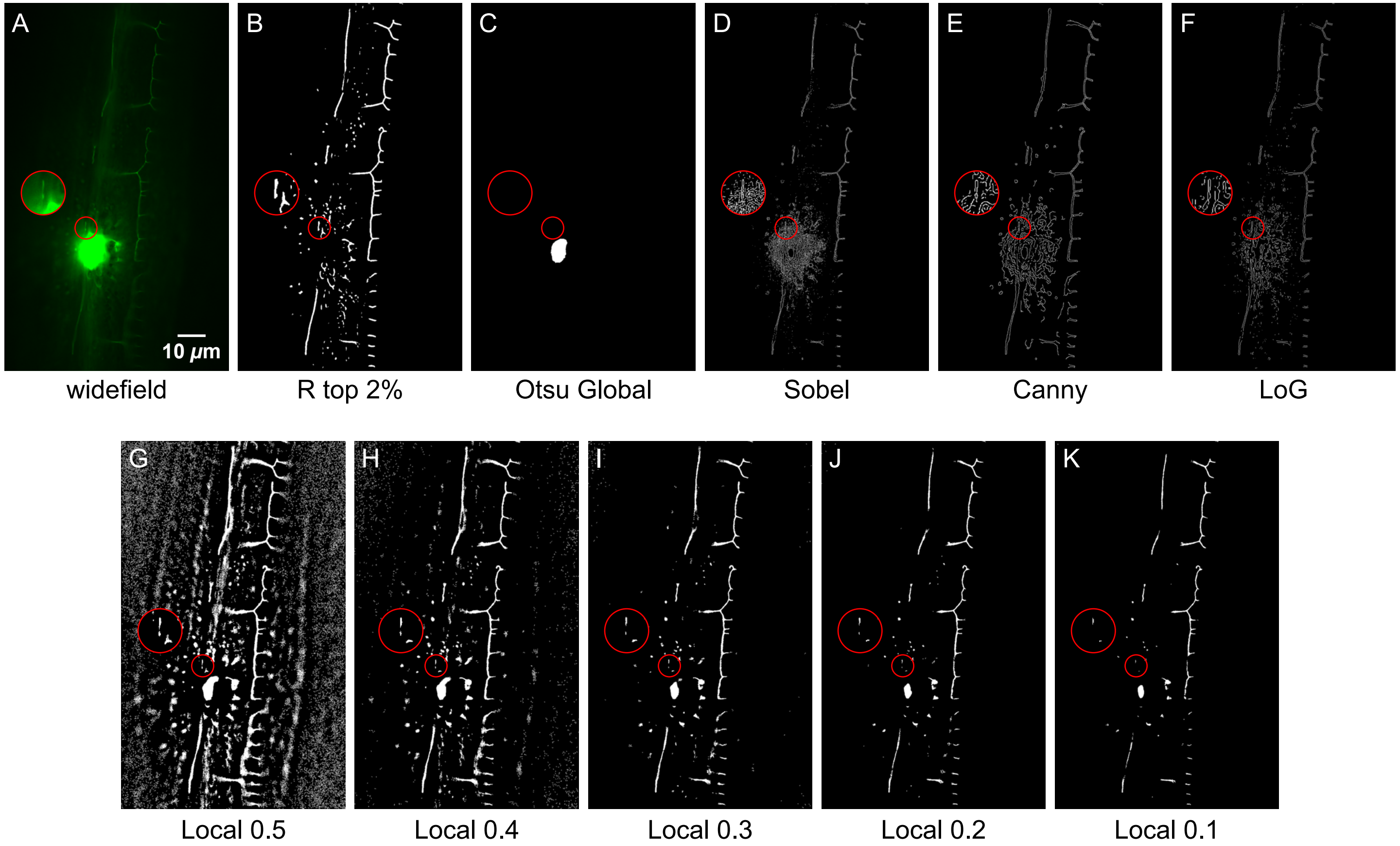


**Fig. S3. Different image processing method for fiber detection**

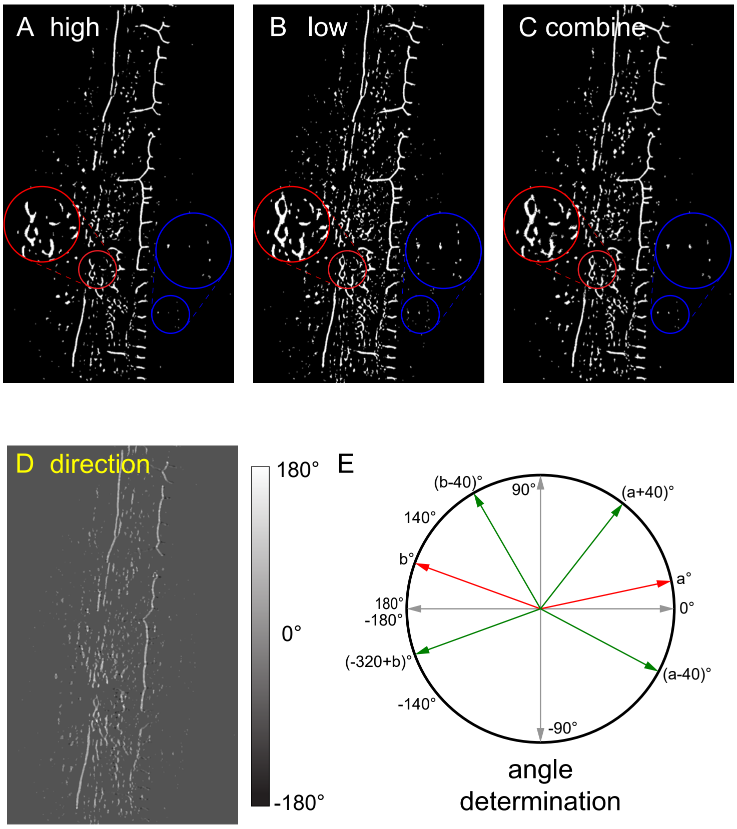


**Fig. S4. directionality consideration**

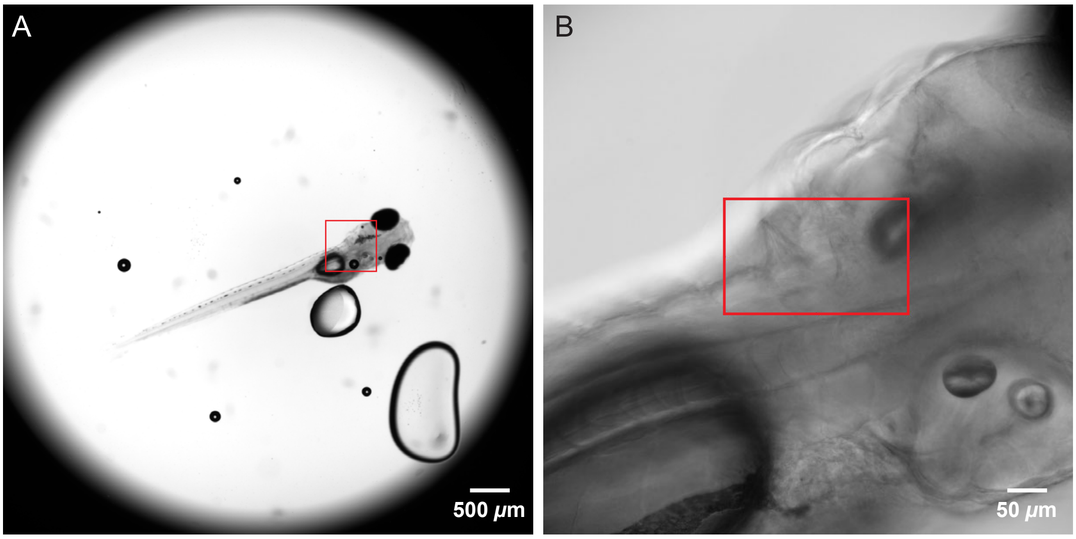


**Fig. S5. zebrafish imaging area**
